# Meta-analysis of transcriptomes of SARS-Cov2 infected human lung epithelial cells identifies transmembrane serine proteases co-expressed with ACE2 and biological processes related to viral entry, immunity, inflammation and cellular stress

**DOI:** 10.1101/2020.05.12.091314

**Authors:** Wasco Wruck, James Adjaye

## Abstract

The COVID-19 pandemic resulting from the severe acute respiratory syndrome coronavirus 2 (SARS-CoV-2) which emerged in December 2019 in the Chinese city of Wuhan in the province Hubei has placed immense burden on national economies and global health. At present neither vaccination nor therapies are available although several antiviral agents such as remdesivir, originally an Ebola drug, nelfinavir, an HIV-1 protease inhibitor and other drugs such as lopinavir have been evaluated. Here, we performed a meta-analysis of RNA-sequencing data from three studies employing human lung epithelial cells. Of these one focused on lung epithelial cells infected with SARS-CoV-2. We aimed at identifying genes co-expressed with angiotensin I converting enzyme 2 (ACE2) the human cell entry receptor of SARS-CoV-2, and unveiled several genes correlated or inversely correlated with high significance, among the most significant of these was the transmembrane serine protease 4 (TMPRSS4). Serine proteases are known to be involved in the infection process by priming the virus spike protein. Pathway analysis revealed papilloma virus infection amongst the most significantly correlated pathways. Gene Ontologies revealed regulation of viral life cycle, immune responses, pro-inflammatory responses-several interleukins such as IL6, IL1, IL20 and IL33, IFI16 regulating the interferon response to a virus, chemo-attraction of macrophages, last and not least cellular stress resulting from activated Reactive Oxygen Species. We believe that this dataset will aid in a better understanding of the molecular mechanism(s) underlying COVID-19.

## Introduction

Severe acute respiratory disease COVID-19 is a result of infections with the coronavirus SARS-CoV-2 first reported in the Chinese city Wuhan, Province Hubei, in December 2019 and has since 11 March 2020 been designated as a pandemic by WHO. The origin of the virus is still under investigation although some studies suggest that it was transferred from another species, possibly pangolins^1^ or bats ^2^. At the end of April 2020 the number of globally confirmed cases of COVID-19 exceeded 3 million and recorded deaths beyond 200,000 in the real-time statistics of the Johns Hopkins University^3^. Due to many unreported and asymptomatic cases, the infection fatality rate (IFR) is difficult to determine however Verity *et al.* estimate approximately 0.66% (0·39–1·33) in China^4^. The age-associated IFR is approximately 7.8% for those above 80 years^4^. Drugs have been repurposed for stabilizing COVID-19, but these are not effective therapies^5^, examples include such as remdesivir (Ebola)^6,7^, hydroxy-chloroquine (Malaria)^6,7^ and nelfinavir (HIV)^8^. Another treatment option is to indirectly immunize individuals with plasma from convalescent COVID-19 patients^9^. Further approaches aim at mimicking the human virus cell entry receptor ACE2^2^ with human recombinant soluble ACE2 (hrsACE2)^10^. The cell entry receptor ACE2 associates with transmembrane proteases which prime the spike protein of the virus. Hoffmann *et al.* assigned this task to the protein TMPRSS2^11^ which they identified in the predecessor virus SARS-CoV-2 and assume it is the same for SARS-CoV-2. The protease can be inhibited by existing compounds to interrupt further propagation of the virus in the human host.

Here, we describe a meta-analysis focussing on the transcriptome data from human lung epithelial cells including samples infected with SARS-CoV-2 from a study described by Blanco Melo *et al.*^12^. We directed the exploration to co-expression with the known CoV-2 receptor ACE2. The analysis revealed a signature consisting of 72 genes significantly co-expressed with ACE2 either with positive or negative Pearson correlation. Of the transmembrane serine proteases, the most significantly co-expressed with ACE2 was TMPRSS4, suggesting it to be a putative druggable target.

## Results

### Cluster analysis of SARS-CoV-2 infected cells compared to other non-infected lung cells

Figure 1 shows a hierarchical cluster analysis of all samples used in this meta-analysis. SARS004 cells infected with CoV-2 cluster together with mock-infected SARS004 cells but separated from all other lung epithelial cells and lung carcinoma cell lines which together we consider as control in this analysis. The table of Pearson correlation co-efficients (suppl. Table 2) reflects the grouping implicated by the hierarchical clustering: SARS-CoV-2 samples have highest correlation (r=0.9884-0.9936) to the mock-infected SARS cells.

**Figure 1:**
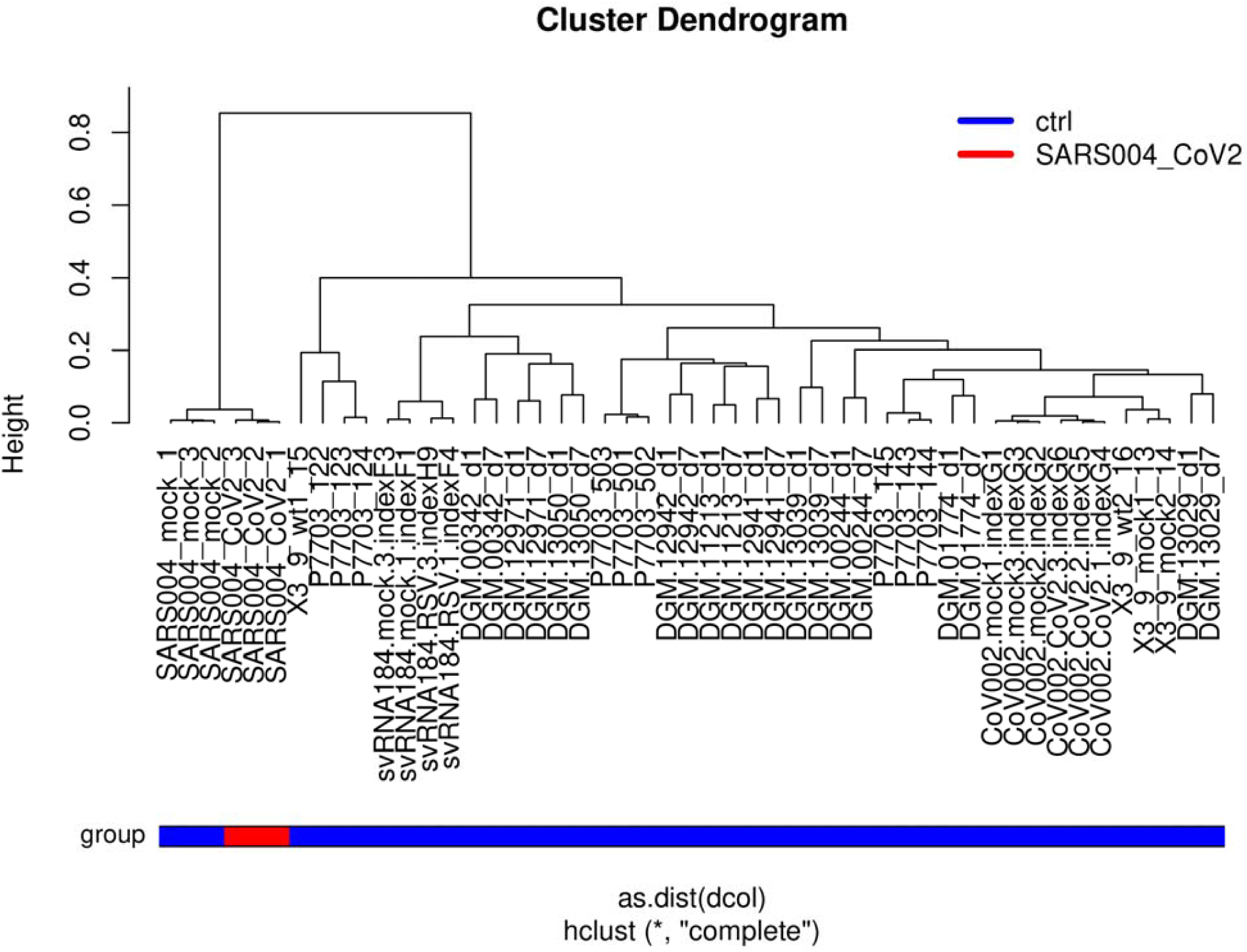
SARS004 cells infected with CoV-2 cluster together with mock-infected SARS004 cells but separated from all other lung epithelial cells.

### Analysis of genes with correlated and anti-correlated expression with ACE2

Building on the knowledge about ACE2 as receptor of the SARS-CoV-2 virus we set out to indentify genes with highly correlated expression with ACE2 with the aim of elucidating the molecular mechanisms underlying COVID-19. Figure 2 shows the genes most significantly (Bonferoni-adjusted p < 1E-11) correlated (red to yellow in last column) or anti-correlated (green) with ACE2 (full table in suppl. Table 3). The transmembrane serine protease 4 (TMPRSS4) is one of the most significantly correlated (r=0.9142, p=4.59E-20) with ACE2 therefore implying a major role in priming the CoV-2 spike protein. CXCL17 (r=0.9273, p=1.1E-21), ABCA12 (r=0.9256, p=1-92E-21) and ATP10B (r=0.9193, p=1.14E-20) had marginally higher correlation with ACE2 while another transmembrane protease TMPRS11E (r=0.9121, p=7.91E-20) had a slightly lower correlation. The expression of CXCL17 is probably due to a reaction on the infection by chemo-attracting macrophages^13^. A The role of ATP binding cassette subfamily A member 12 (ABCA12) is not fully elucidated with respect to COVID-19 but assumed to transport lipid via lipid granules to the intracellular space and transporting specific proteases – in the case of harlequin ichtyosis associated with desquamation^14^. Current knowledge on ATP10B is scarce. However, Wilk *et al.* (Table 3 in their publication) report Atp10b gene expression levels as highly inversely (negative) correlated with influenza gene expression changes in infected C57BL/6J mice^15^. In Figure 3a a cluster analysis and gene expression heatmap of the 72 most significantly (Bonferoni-adjusted p < 1E-11) correlated and anti-correlated genes with ACE2 shows close clustering of the serine protease TMPRSS4 with ACE2. Also in this analysis of 72 genes SARS-CoV-2 (red color bar) cluster together with mock-infected SARS lung epithelial cells separated from the other lung cells (blue color bar indicates control). In the heatmap presented in Figure 3b, TMPRSS family members TMPRSS11D/E and TMPRSS4 show close clustering with ACE2 but also TMPRSS2 and TMPRSS13 have similar expression in all experiments, especially in the SARS-CoV-2 infected samples.

**Table 1:** Datasets used in this focussed meta-analysis.

| dataset | description | reference |
| --- | --- | --- |
| GSE147507 | primary human bronchial epithelial cells and lung adenocarcinoma infected with CoV2 or Mock | Blanco-Melo et al. Biorxiv 2020 |
| GSE146482 | human bronchial epithelium cell line BEAS-2B | Mukherjee SP et al. unpublished |
| GSE85121 | small airway epithelium brushing | Staudt MR et al. Respir Res 2018 May 14;19(1):78. PMID: 29754582 |

**Table 2:**
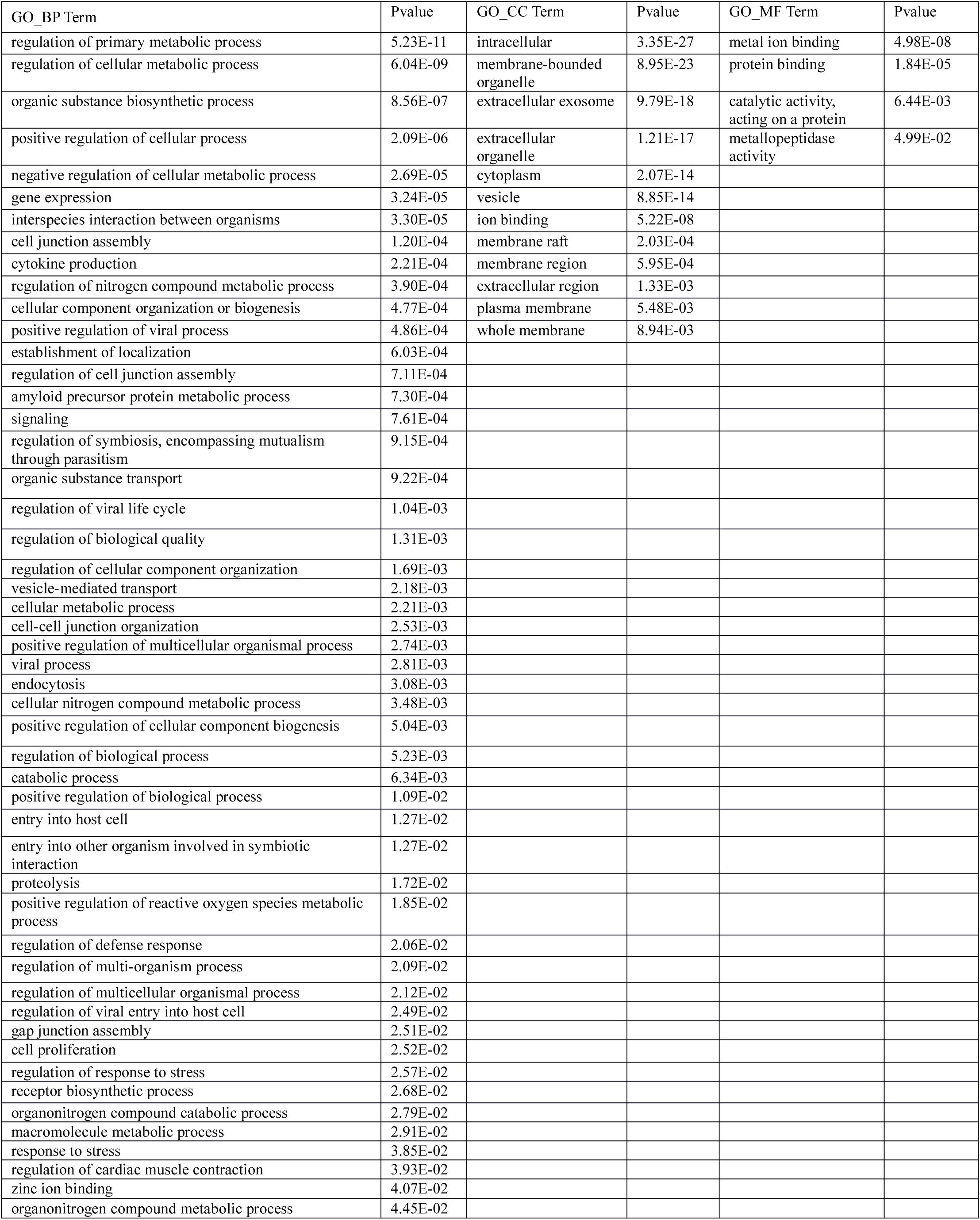
Selected over-represented GOs in genes significantly correlated with ACE2.

**Table 3:** GOs (all Biological Process) related to immune system in genes correlated with *ACE2*.

| Term | Pvalue |
| --- | --- |
| myeloid cell activation involved in immune response | 5.397772E-07 |
| cell activation involved in immune response | 4.365561E-05 |
| negative regulation of immune system process | 0.0033136313 |
| immune system process | 0.0048604032 |
| regulation of chemokine production | 0.0048612934 |
| regulation of interleukin-1 production | 0.0048612934 |
| positive regulation of innate immune response | 0.0078573986 |
| immune response | 0.008554933 |
| positive regulation of myeloid leukocyte cytokine production involved in immune response | 0.0180984932 |
| innate immune response-activating signal transduction | 0.0219067163 |
| positive regulation of T cell activation | 0.0220941875 |
| astrocyte activation involved in immune response | 0.0276437419 |
| microglial cell activation involved in immune response | 0.0276437419 |
| immune response-regulating signaling pathway | 0.0286071006 |
| negative regulation of immune effector process | 0.0375778444 |

**Figure 2:**
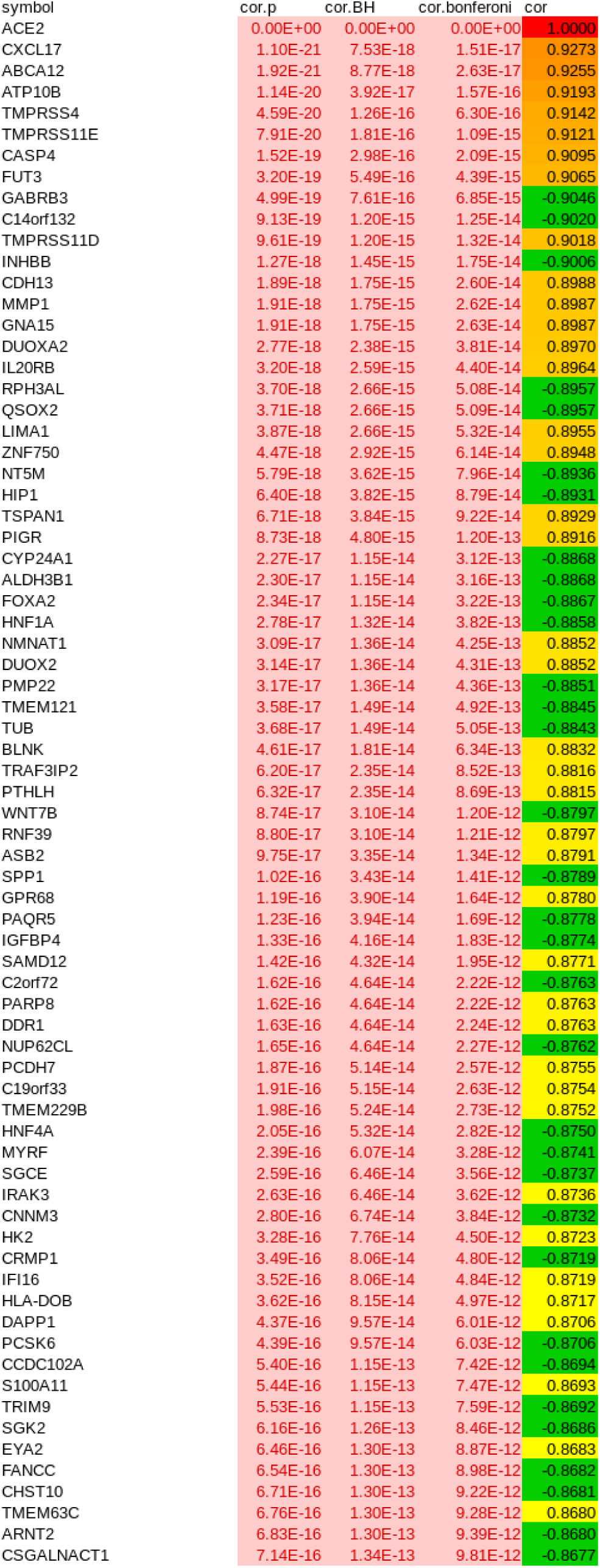
Most significantly (Bonferoni-adjusted p < 1E-11) correlated (red to yellow in last column) or anti-correlated (green) genes with ACE2. The transmembrane serine protease 4 (TMPRSS4) is one of the most significantly correlated (r=0.9142, p=4.59E-20) with ACE2 suggesting a major role in priming the CoV2 spike protein. CXCL17 (r=0.9273, p=1.1E-21), ABCA12 (r=0.9256, p=1-92E-21) and ATP10B (r=0.9193, p=1.14E-20) had marginally higher correlation with ACE2 while another transmembrane protease TMPRS11E (r=0.9121, p=7.91E-20) had slightly lower correlation.

**Figure 3:**
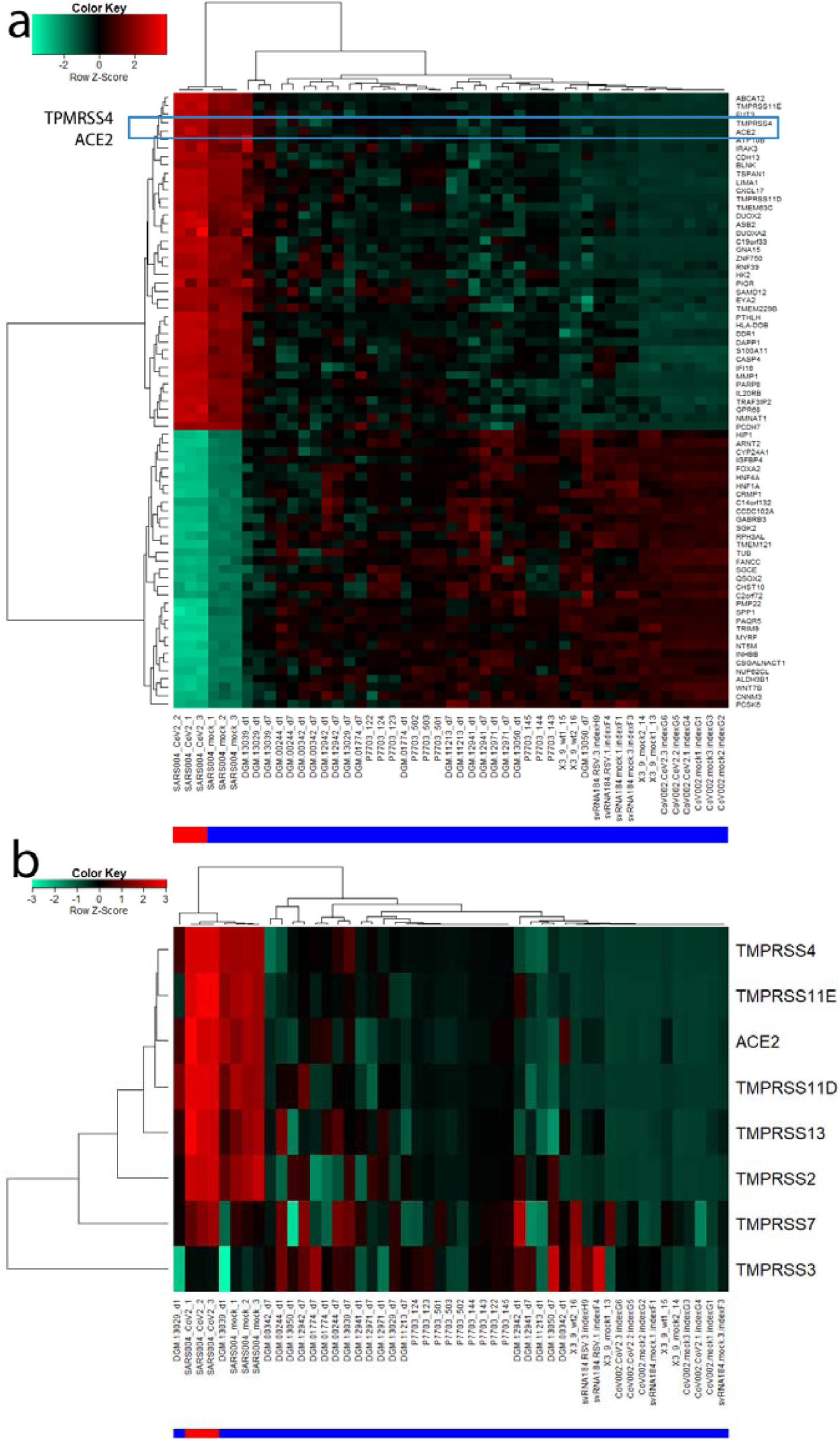
(a) Cluster analysis and gene expression heatmap of 72 most significantly (Bonferoni-adjusted p < 1E-11) correlated and anti.correlated genes with ACE2 shows close clustering of the serine protease TMPRSS4 with ACE2. (b) Heatmap of TMPRSS family members shows close clustering of TMPRSS11D/E and TMPRSS4 with ACE2 but also TMPRSS2 and TMPRSS13 have similar expression, particularly in SARS-CoV-2 infected samples. The red color bar indicates SARS-CoV-2, blue color bar control.

### Pathway analysis of genes co-regulated with ACE2

In order to investigate the functionality of genes interacting with *ACE2* we filtered genes correlated with ACE2 by Bonferoni-adjusted p-value < 0.05 and Pearson correlation coefficient > 0.6. 1891 genes fulfilled these criteria and were subjected to over-representation analysis of KEGG pathways^16^. The most significantly overrepresented pathways associated with the 1891 genes correlated with *ACE2* (Figure 4a, suppl. Table 4) are for example, *Bacterial invasion of epithelial cells (q=4.4E-06), Human papillomavirus infection (q=0.0006), Transcriptional misregulation in cancer (q=0.0006)* and *Endocytosis (q=0.002)*. This reflects the mechanisms of virus infection via invasion of epithelial cells and endocytosis.

**Figure 4:**
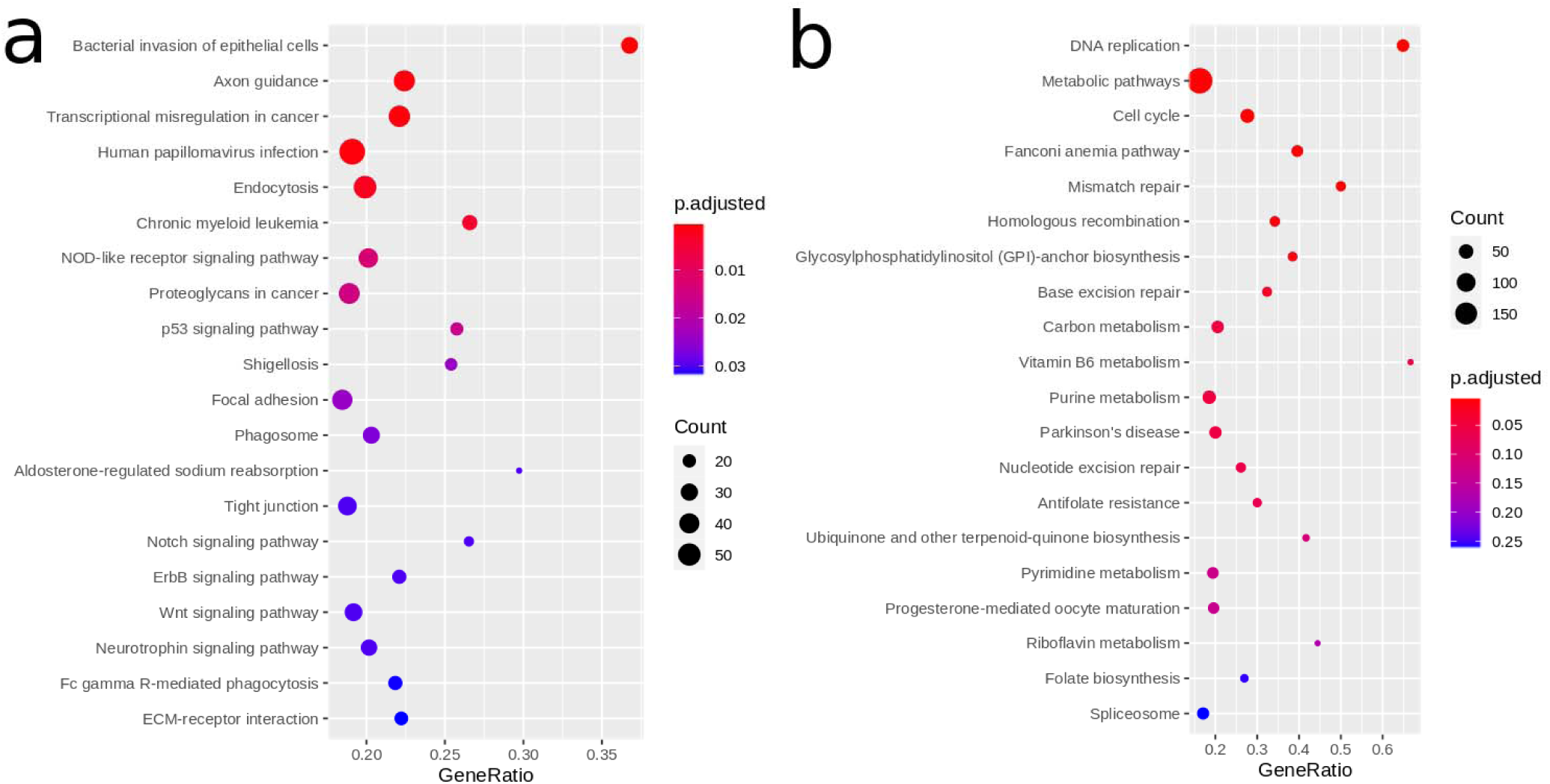
(a) The five most significantly overrepresented pathways correlated with ACE2 are *Human papillomavirus infection, Bacterial invasion of epithelial cells, Endocytosis, Axon Guidance* and *Transcriptional mis-regulation in cancer*. (b) The six most significantly overrepresented pathways anti-correlated with ACE2 are *DNA replication, Metabolic pathways, Cell cycle, Fanconi anemia pathway, Mismatch repair* and *Homologous recombination*. Many of these pathways are associated with DNA processing or repair. That these are down-regulated upon infection with CoV-2 is in line with reports about interferon and interferon stimulated genes (ISGs) inhibiting virus replication^16^. This would be a defense against the attempts of the virus to recruit the host’s DNA repair and homologous recombination mechanisms as Gillespie et al. report^18^.

### Pathway analysis of genes anti-correlated with ACE2

Analogously to the positively correlated genes we also examined the negatively correlated genes with ACE2 by filtering for Bonferoni-adjusted p-value < 0.05 and Pearson correlation coefficient < −0.6. 1993 genes passed these filtering criteria and were subjected to over-representation analysis of KEGG pathways^15^. The most significantly over-represented pathways in the 1993 genes negatively correlated with *ACE2* (Figure 4b, suppl. Table 5) are *DNA replication (q=1E-12), Metabolic pathways (q=1.86E.08), Cell cycle (q=1.1E-05), Fanconi anemia pathway (q=1.24E-05), Mismatch repair (q=9.89E-05)* and *Homologouos recombination (q=0.0046)*. Many of these pathways are associated with DNA processing or repair. That these are over-represented in genes down-regulated upon infection with CoV-2 is in line with reports about interferon and interferon stimulated genes (ISGs) inhibiting virus replication^17^. This would be a defense against the attempts of the virus to recruit the host’s DNA repair and homologous recombination mechanisms ^18^.

### GO analysis of genes co-regulated with ACE2

We furthermore assessed the GOs over-represented in the 1891 genes positively and the 1993 genes negatively correlated with *ACE2*. Table 2 shows a selection of significant GOs from all three categories Biological Process (BP, Figure 5a), Cellular Component (CC) and Molecular Function (MF), suppl. Table 6 provides the full table and suppl. Table 7 provides the full table for the 1993 genes negatively correlated with *ACE2*. Amongst the GO-BPs, metabolic processes are the most significant. GO-BP terms such as *Interspecies interaction between organisms, Cytokine production* and *positive regulation of viral process* reflect activated mechanisms post-viral infection. Interestingly, we found *regulation of coagulation* amongst the GO-BPs what may help elucidate reports about co-agulation in acro-ischemic COVID-19 patients^19^. In the GO-CCs, the terms *intracellular* and *membrane-bounded organelle* are most significant. In the GO-MFs, the terms *metal ion binding* and *protein binding* emerge as most significant due probably reflecting the binding of the virus proteins to the host cells. For the full gene lists associated with these terms refer to suppl. Tables 5 and 6.

**Figure 5:**
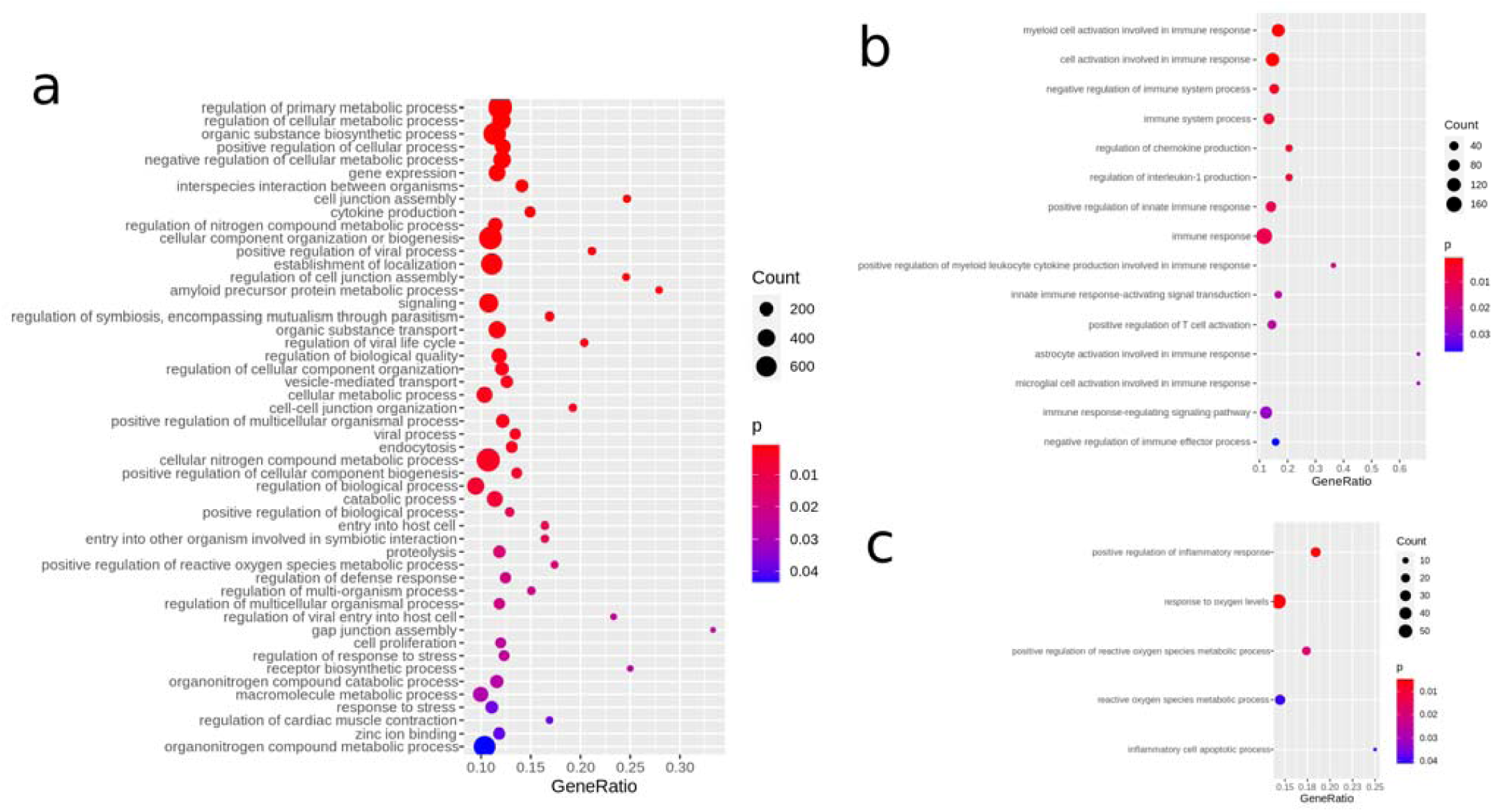
GO analysis reflects virus entry and immune response involving ROS and inflammation. (a) Selected GOs (Biological processes) shows interaction of virus and host, cell-junction organization, endocytosis, reaction involving cytokine production. (b) Immunity-related GOs illustrate the immune response involving activation of myeloid cells and T-cells and interleukin-1 and other chemokine production. (c) GOs associated with ROS and inflammation demonstrate involvement of ROS and inflammation leading to apoptosis.

### Immune system associated GOs co-regulated with ACE2

Table 3 and Figure 5b show GOs (all Biological Processes) related to the immune system over-represented in genes correlated with *ACE2*. Myeloid cells involved in the immune response (GO:0002275, p=5E-07) as well as T-cells (GO:0050870, p=0.02) are activated. Chemokines, in particular interleukin-1 are produced (GO:0032642, p=0.0049, GO:0032652, p=0.0049) also IL33 and TNF. Additionally, positive regulation of the innate immune response was prominent (GO:0045089, p=0.0079). *CXCL17* was the gene with the highest correlation with *ACE2* and is involved in immune system process - as described above by chemo-attracting macrophages^13^. Among the most significantly *ACE2*-correlated genes was *IFI16* which is associated with several immune system GOs and known as regulator of the interferon response to viruses^20^ and will be described in more detail in the next section about *Protein interaction networks*. Also in the protein interaction network of the most significantly ACE2-correlated genes was the interleukin 20 receptor B (IL20RB) which appeared in several GOs listed in Table 3. With respect to viruses, there is meagre knowledge on IL20RB, however a study reported over-expression in the pneumonia causing avian influenza A H7N9 virus^21^.

### GOs associated with inflammation and reactive oxygen species (ROS) amongst genes co-regulated with ACE2

Table 4 and Figure 5c show GOs (all Biological Processes) related to inflammation and ROS over-represented in genes correlated with ACE2. The GO, *positive regulation of inflammatory response* (p=0.0039) would imply that an inflammatory process is induced which finally leads to apoptosis - as demonstrated by the GO *inflammatory cell apoptotic process* (p=0.042). The virus receptor ACE2 is involved in the *positive regulation of reactive oxygen species metabolic process* (p=0.0185). Induction of ROS by a respiratory virus and subsequent inflammation has been reported Jamaluddin *et al.*^22^.

**Table 4:** GOs (all Biological Process) related to inflammation and reactive oxygen species (ROS) in genes correlated with *ACE2*.

| Term | Pvalue |
| --- | --- |
| positive regulation of inflammatory response | 0.0039262994 |
| response to oxygen levels | 0.0043898848 |
| positive regulation of reactive oxygen species metabolic process | 0.0185189125 |
| reactive oxygen species metabolic process | 0.0408852411 |
| inflammatory cell apoptotic process | 0.0420672178 |

### Protein interaction networks

We further restricted the set of ACE2-correlated or -anti-correlated genes by drastically filtering with Bonferoni-adjusted p < 1E-11 in order to construct a human readable protein interaction network of the most significant proteins (Figure 6). The protein interaction network generated from correlated and anti-correlated genes with *ACE2* shows *IFI16 (r=0.8719), LIMA1 (r=0.8955), CNNM3 (r=-0.8732), HNF4A (r=-0.8750), TRAF3IP2 (r=0.8816), ASB2 (r=0.8791)* and *FANCC* (r=-0.8682) as hub genes (interactors from the BioGrid database are marked in red, original *ACE2*-correlated/anti-correlated genes are marked in green, hub genes and *ACE2* are highlighted with yellow shading). Interferon plays a major role in the host response to a virus and Thompson *et al.* reported that IFI16 – one of the hub genes in our network - controls the interferon response to DNA and RNA viruses^20^. Lin and Richardson review that *LIMA1* (formerly EPLIN) – which is known to enhance bundling of actin filaments^23^ - mediates the interaction between Cadherins and Actin in the context of adherens junctions - playing a role in measles virus infection - via trans-binding with molecular interactors on adjacent cells^24^. In line with that LIMA1 is connected with E-cadherin (CDH1) in the network and also cadherin 13 (CDH13) is part of it and among the most significantly correlated genes to ACE2. The connection of ACE2 to calmodulin 1 (CALM1) is based on a publication by Lambert *et al.*^25^ in which they show that CALM1 interacts with the corona virus receptor ACE2 and inhibits shedding of its ectodomain^25^. CALM1 inhibitors in turn can revert this process so that the ACE2 ectodomain is shed partially mediated by a metalloproteinase^25^. The direct connection between CALM1 and LIMA1 was found in a large-scale interactome study^26^. The involvement of the hub gene Fanconi anemia complementation group C (FANCC), although not experimentally proven, might reflect recruitment of DNA repair and homologous recombination mechanisms from the host by the virus.

**Figure 6:**
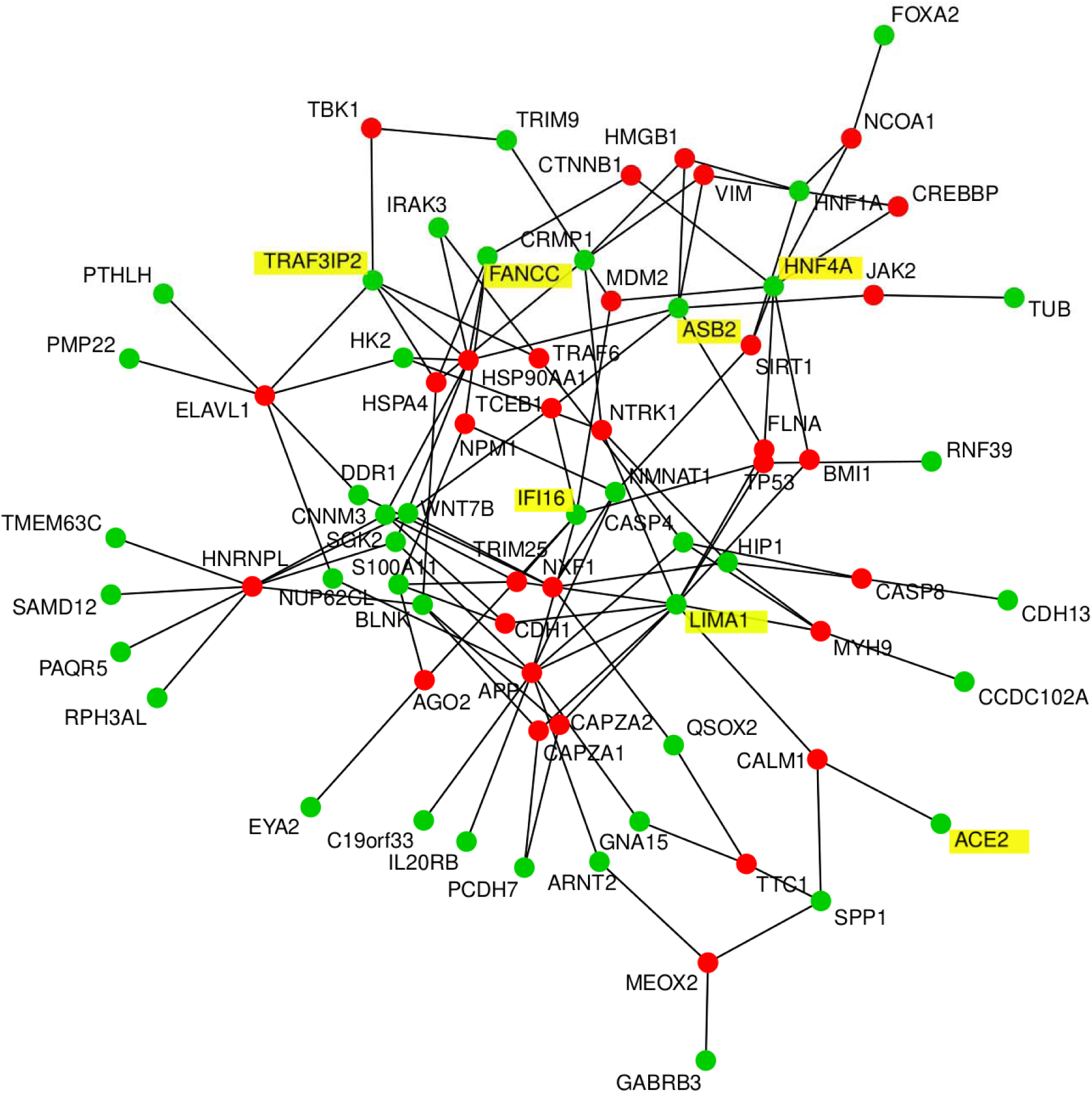
Protein interaction network of genes most significantly (Bonferoni-adjusted p < 1E-11) correlated and anti-correlated genes with *ACE2* shows *IFI16 (r=0.8719), LIMA1 (r=0.8955), CNNM3 (r=-0.8732), HNF4A (r=-0.8750), TRAF3IP2 (r=0.8816), ASB2 (r=0.8791)* and *FANCC* (r=-0.8682) as hub genes. Genes found as interactors in the BioGrid database are marked in red, the original geneset of *ACE2*-correlated genes is marked in green, hub genes and *ACE2* have yellow shading.

### Role of TMPRSS4 and other TMPRSS gene family members

A pivotal result of this meta-analysis is the transmembrane serine protease TMPRSS4 which was one of the most significantly correlated genes with ACE2. Additionally TMPRSS11E (r=0.9121, Bonferoni-corrected p=1.09E-15) and TMPRS11D (r=0.9018, Bonferoni-corrected p=1.32E-14) from the same gene family were found more significant than TMPRSS2 which however was still significantly correlated with ACE2 (r=0.767, Bonferoni-corrected p=1.79E-6). The assignment of the SARS-CoV-2 spike protein priming functionality to TMPRSS2 was based on the assumption that it would be the same as for its predecessor SARS-CoV ^11^. However, this assumption must not necessarily hold true. See the heatmap in Figure 3b which illustrates expression of transmembrane serine proteases. The highly significant co-expression of TMPRSS4 with ACE2 and its relevant role as transmembrane serine protease has enabled us to hypothesize that TMPRSS4 might also be involved in priming the SARS-CoV-2 spike protein. We therefore anticipate that inhibitors of TMPRSS4, TMPRS11D and TMPRS11E - besides those for TMPRSS2 - could be a promising subject of further research.

## Discussion

In this meta-analysis we compared RNA-seq data of lung cells infected with SARS-CoV-2 and other lung cells with particular focus on correlated gene expression with the known SARS-CoV-2 receptor gene *ACE2*. We found a signature of genes positively or negatively correlated with *ACE2* amongst which the most outstanding was the transmembrane serine protease *TMPRSS4*. In a recent publication Hoffmann *et al*^11^. inferred from the knowledge that the preceding virus SARS-CoV uses ACE2 as receptor for entry and the serine protease TMPRSS2 for spike protein priming that the new virus SARS-CoV-2 ^11^ would do the same. While the involvement of the receptor ACE2 appears to be established^1,10^ the use of TMPRSS2 for spike protein priming appears not fully settled as Hoffmann *et al.* still have to concede “that SARS-CoV-2 infection of Calu-3 cells was inhibited but not abrogated by camostat mesylate” (serine protease inhibitor with activity against TMPRSS2)^11^. The high significance (r=0.9142, p=4.59E-20) in our co-expression analysis with ACE2 suggests that TMPRSS4 is a considerable candidate for spike protein priming. However, TMPRSS4 is closely related to TMPRSS2 which both can proteolytically cleave the hemagglutinin of influenza viruses^27^. Further transmembrane serine proteases TMPRS11D (r=0.9018, p=9.61E-19) and TMPRS11E (r=0.9121, p=7.91E-20) in our analysis also emerged more significant than TMPRSS2 (r=0.767, p=1.3E-10). The TMPRS11 family member TMPRS11A was found to enhance viral infection with the first coronavirus SARS-CoV by spike protein cleavage in the airway^28^. Thus, we should not exclude the probability that other members of the TMPRSS gene family may be proteases for the SARS-CoV2 spike protein. TMPRSS2 inhibitors have been proposed by Hoffmann *et al.*^11^ - and earlier for the SARS-Cov virus by Kawase *et al.*^29^ as working best together with cathepsin B/L inhibitors. We propose to investigate the effect of TMPRSS4 inhibitors in further research. As TMPRSS4 has been implicated in the invasion and metastasis of several cancers it has been also considered as target for cancer therapy for which a modest inhibitory effect of the above mentioned inhibitors in TMPRSS4-overexpressing SW480 cells was reported ^30^. Interestingly, also *TMPRSS2* is connected to epithelial carcinogenesis as consistently overexpressed in prostate cancer ^31^, and later a gene fusion of *TMPRSS2* and *ERG* was reported as the predominant molecular subtype of prostate cancer ^32^, where *TMPRSS2* however only contributes untranslated sequence^33^. Assuming that co-expression with *ACE2* is an indication that TMPRSS4 may prime the SARS-CoV2 spike protein we suggest that further development and testing of more effective TMPRSS4 inhibitors in *in vitro* and *in vivo* models could support the translation into clinical settings. However, we have to state the limitation that this study is a meta-analysis based solely on transcriptome and not proteome data.

Besides the identification of *TMPRSS4*, we found several significantly over-represented GOs and pathways such as Endocytosis, Papilloma virus infection and Bacterial invasion of epithelial cells for which we provide full gene lists to foster further elucidation of disease mechanisms. Genes from the constructed protein interaction network provide a first snapshot of a comprehensive image: *IFI16* controls the interferon response to the virus^20^, *LIMA1* mediating the interaction between Cadherins (*CDH1, CDH13*) and Actin in the context of adherens junctions potentially playing a role in virus infection, *CALM1* inhibits shedding of the ectodomain of the virus receptor *ACE2* ^25^.

Furthermore, GO analyses revealed several biological processes related to viral cell entry, host reaction, immune response, ROS, inflammation and apoptosis. This led us to propose a cascade of events taking place post SARS-CoV-2 entry into host cells-illustrated in Figure 7 together with possible drug targets. The coronavirus SARS-CoV-2 docks at the receptor ACE2 on the membrane of the human epithelial cell, the early stage of infection. According to reports by Monteil *et al.* these processes can be blocked with recombinant hrsACE2^10^. Transmembrane serine proteases TMPRSS2 mediate SARS-CoV-2 cell entry via ACE2^11,34^. TMPRSS2 has been described as a mediator of ACE2-coupled endocytosis in the first SARS-CoV ^35^ and by previous publication also for SARS-CoV2^11^. However, we identified high levels of co-expression between ACE2 and TMPRSS4 and other members of the TMPRSS family and hypothesize that any of these additional family members might have the same function as the well described TMPRSS2. As a consequence, we propose that besides TMPRSS2 also other TMPRSS family members could be targets of pharmaceutical intervention warranting further research. After entering the cell, SARS-CoV-2 RNA is released, replicated and packaged again. Drugs can target the viral protease and the polymerase needed for replication^36^. Replication can further be inhibited by interferon and interferon-stimulated genes (ISG)^17^which we also found evidence for in negatively correlated replication pathways (e.g. DNA replication and homologous recombination). This depends on a healthy immune response and may be impaired in individuals with a weak immune system due to age or underlying diseases. The virus is packaged and released into the extracellular space where it can be attacked by macrophages chemo-attracted by CXCL17^13^. Also T-Cells can be involved in the immune response. We found evidence for their activation in GO analysis by associated interleukins IL1 and IL7. It is tempting to speculate that the severity of the clinical manifestations such as the acute respiratory failure and also failure in other organs depend on the state of the immune system which decreases with age or diseases such as diabetes. The involvement of ACE2 in the renin-angiotensin system as antagonist of ACE in regulating blood pressure via Angiotensin II, vasoconstriction, dilation and its protective role against lung injury ^37^ are additional factors which correlate with age. This is confirmed by reports from Wadmann *et al.*^38^ about Centers for Disease Control and Prevention (CDC) data from 14 U.S. states that 50% hospitalized COVID-19 patients had pre-existing high blood pressure^38^. In their study about ACE2 in the preceeding SARS-CoV virus Imai *et al.* ^37^ found that *ACE2* protects against lung injury and acid-induced lung injury in a *Ace2*-knockout mouse can be improved by an inhibitor of the Angiotensin II receptor AT1^37^. The results of clinical studies but also statistics on hypertension and even more important statistics on pharmacological treatment of hypertension in COVID-19 patients may shed light on the discussions if treatment with ACE inhibitors and angiotensin receptor blockers (ARBs) are detrimental^39^ or beneficial^40^.

**Figure 7:**
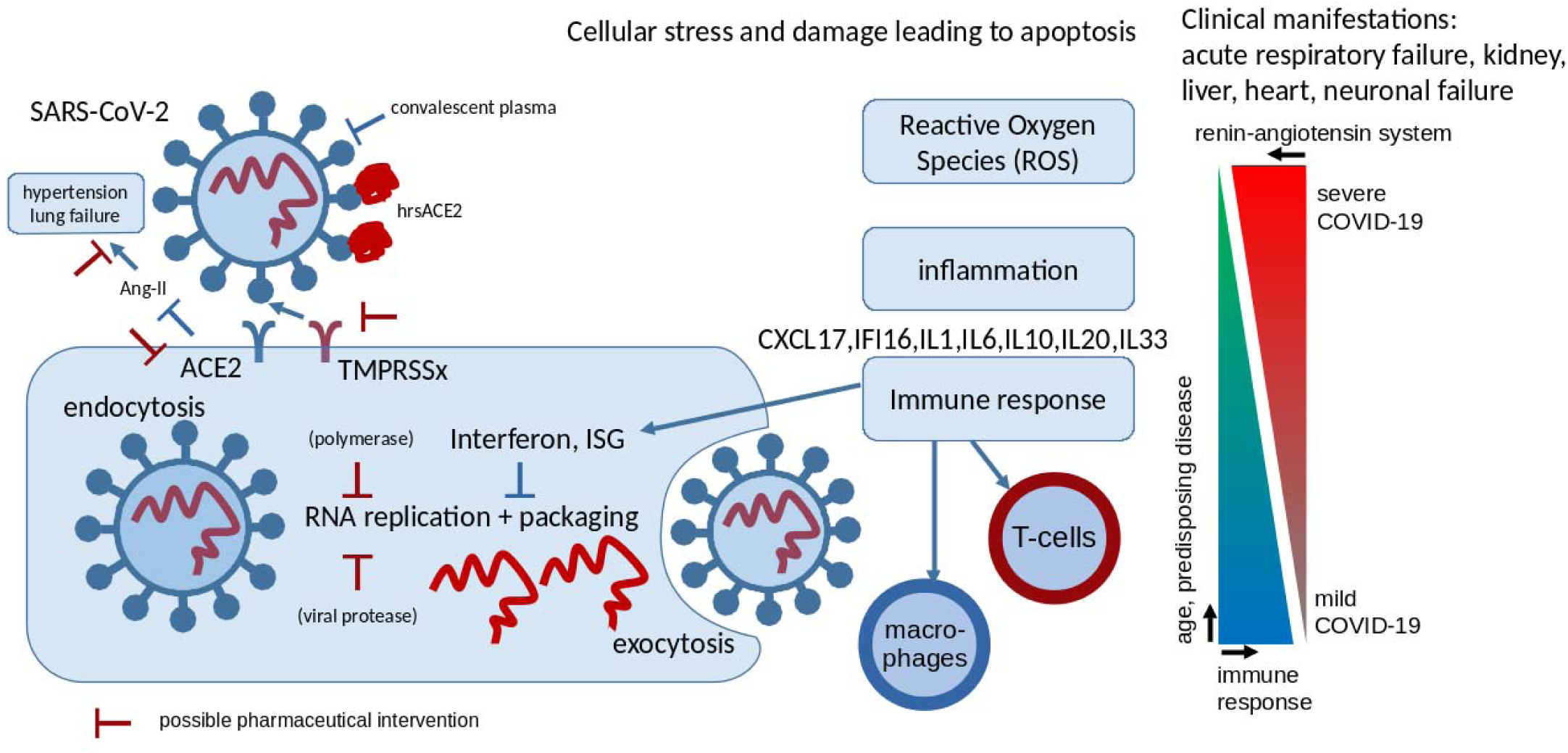
Scheme of SARS-CoV-2 infection. The coronavirus SARS-CoV-2 docks at the receptor ACE2 on the membrane of the human epithelial cell. Transmembrane serine proteases TMPRSSx mediate SARS-CoV-2 cell entry via ACE2. TMPRSS2 was reported for this in the first SARS-CoV and by previous publication also for SARS-CoV-2 but we hypothesize that due to co-expression with ACE2, TMPRSS4 and other members of the TMPRSS family may well perform this task. We suggest that inhibitors of TMPRSS4 and other TMPRSS family members might have therapeutic potential. Upon entry into the cell, viral RNA is released, replicated and packaged again. Replication can be inhibited by interferon and interferon stimulated genes (ISG) what we also saw in negatively correlated replication pathways (e.g. DNA replication and homologous recombination). This indicates a healthy immune response and may be impaired in persons with a weak immune system due to age or disease. The packaged virus is released from the cell and can be attacked by macrophages chemo-attracted by CXCL17 - or T-cells for which we found evidence for by GO analysis and by associated interleukins IL1 and IL7. It is tempting to speculate that the severity of the clinical manifestations such as the acute respiratory failure and also failure in other organs depends on the quality of the immune system decreasing with age or diseases such as diabetes. The involvement of ACE2 in the renin-angiotensin system as antagonist of ACE in regulating blood pressure via Angiotensin II (Ang-II), vasoconstriction, dilation and its protective role against lung injury are additional factors which correlate with age.

We conclude, that our meta-analysis of RNA-Seq data of lung cells partially infected with SARS-CoV-2 identified the transmembrane serine protease *TMPRSS4* as one of the most significantly correlated genes with the virus receptor *ACE2*. We propose that inhibitors of TMPRSS family members TMPRSS4, TMPRSS11D and TMPRSS11E besides TMPRSS2 are worthwhile testing i*n vitro* and if that turns out to be successful and toxicity can be excluded potentially for i*n vivo* studies. As clinicians, pathologists and scientists are struggling to get to grips with and understanding of the damage wrought by SARS-CoV-2 as it invades the body, we hope that our analyses and dataset will contribute to a better understanding of the molecular basis of COVID-19.

## Methods

### Sample collection of lung cell RNA-Seq data

Next-generation sequencing datasets measured in RNA-Seq experiments with lung cells (GSE147507: Illumina NextSeq 500; GSE146482: Illumina NovaSeq 6000; GSE85121: Illumina HiSeq 2500) were downloaded from NCBI GEO (Table 1, suppl. Table 1). These datasets were provided along with studies by Blanco-Melo *et al.*^12^ (GSE147507) and Staudt e*t al.*^41^ (GSE85121). A final publication related to the dataset GSE146482 is yet to materialize. From accession no. GSE85121 only small airway epithelium brushing cells were used but alveolar macrophages were excluded while from accession no. GSE147507 only human epithelial and adenocarcinoma lung cells were used but ferret cells and updates after March 24 were excluded and from accession no. GSE146482 only control epithelial cells were used but graphene oxide treated cells were excluded. After exclusion of the above mentioned datasets not fitting the target cell type, 49 samples remained useful.

### Data normalization and analysis

After the excluded samples had been filtered from the downloaded RNA-Seq data data was imported into the R/Bioconductor environment^42^. Read counts from accesion nos. GSE147507 and GSE146482 were converted to FPKM (fragments per kilobase of exon model per million reads mapped) using trancript lengths downloaded from ENSEMBL (version GRCh38, p13). Batch effects were removed with the package sva^43^ employing method ComBat^44^. Normalization was performed via the voom method^45^. Pearson correlation coefficients between samples were calculated with the R-builtin method *cor*. Dendrograms were drawn employing the dendextend package^46^ filtering genes for high coefficient of variation above the 75% quantile.

### Detection of genes correlated with ACE2

The Pearson correlation of the normalized gene expression values for all samples was calculated between the gene *ACE2* and each other gene. The method cor.test was applied to calculate the p-value for the t-test for Pearson correlation. The p-value was Bonferoni-corrected by division by the number of genes and additionally adjusted via the Benjamini-Hochberg method^47^. Correlated genes were filtered with a very restrictive criterion (Bonferoni-adjusted-p < 1E-11) - in order to get human readable numbers of genes for heatmap and network generation- and conventional criteria (r > 0.6, Bonferoni-adjuste-p < 0.05) for positively correlated genes and (r < 0.6, Bonferoni-adjusted-p < 0.05) for negatively correlated genes.

### Pathway and gene ontology (GO) analysis

The R package GOstats^48^ was employed for over-representation analysis of positively and negatively correlated genes with the CoV-2 receptor gene *ACE2*. KEGG pathway annotations which had been downloaded from the KEGG database^16^ in March 2018 were used for testing overrepresentation of the positively and negatively ACE2-correlated genes via the R-builtin hypergeometric test.

Dot plots showing the p-value of the hypergeometric test, the ratio of significant genes to all genes in the pathway and the number of significant genes per pathway were plotted via the package *ggplot2*^49^.

### Protein interaction networks

A human protein interaction network was constructed in a similar manner as we described in our previous publication^50^. However, here only direct interactors and no further interactors of interactors were extracted from the Biogrid database version 3.4.161^51^ using the restrictively filtered (Bonferoni-adjusted-p < 1E-11) genes significantly correlated or anti-correlated with ACE2 gene expression. The network was reduced to the n=30 nodes with most interactions and was plotted via the R package *network* ^52^ showing original genes in green and BioGrid-derived interactors in red.

## Author contributions

WW analysed the data and wrote the manuscript. JA supervised the work, co-wrote the manuscript and gave the final approval.

## Acknowledgments

James Adjaye acknowledges support from the Medical faculty of the Heinrich-Heine University, Duesseldorf.

## Supplementary Material

**Supplementary Table 1 (tableS1.xlsx): Detailed description of all samples used in this meta-analysis**

**Supplementary Table 2 (tableS2.xlsx): Pearson correlation coefficients of gene expression of all samples used in this meta-analysis**

**Supplementary Table 3 (tableS3.xlsx): Pearson correlation coefficients and associated p-values (additionally Benjamini-Hochberg and Bonferoni correction) of genes to expression of ACE2**

**Supplementary Table 4 (tableS4.xlsx): Over-represented KEGG pathways in genes positively correlated with *ACE2* gene expression**

**Supplementary Table 5 (tableS5.xlsx): Over-represented KEGG pathways in genes negatively correlated with *ACE2* gene expression**

**Supplementary Table 6 (tableS6.xlsx): Over-represented GOs in genes positively correlated with *ACE2* gene expression**

**Supplementary Table 7 (tableS7.xlsx): Over-represented GOs in genes negatively correlated with *ACE2* gene expression**

## References

1. Zhang, T., Wu, Q. & Zhang, Z. Probable Pangolin Origin of SARS-CoV-2 Associated with the COVID-19 Outbreak. Curr. Biol. 30, 1346-1351.e2 (2020).

2. Zhou, P. et al. A pneumonia outbreak associated with a new coronavirus of probable bat origin. Nature 579, 270–273 (2020).

3. Dong, E., Du, H. & Gardner, L. An interactive web-based dashboard to track COVID-19 in real time. Lancet Infect. Dis. S1473309920301201 (2020) doi:10.1016/S1473-3099(20)30120-1.

4. Verity, R. et al. Estimates of the severity of coronavirus disease 2019: a model–based analysis. Lancet Infect. Dis. S1473309920302437 (2020) doi:10.1016/S1473-3099(20)30243-7.

5. Wang, Y. et al. Remdesivir in adults with severe COVID-19: a randomised, double-blind, placebo-controlled, multicentre trial. The Lancet 0, (2020).

6. Wang, M. et al. Remdesivir and chloroquine effectively inhibit the recently emerged novel coronavirus (2019-nCoV) in vitro. Cell Res. 30, 269–271 (2020).

7. Lai, C.-C., Shih, T.-P., Ko, W.-C., Tang, H.-J. & Hsueh, P.-R. Severe acute respiratory syndrome coronavirus 2 (SARS-CoV-2) and coronavirus disease-2019 (COVID-19): The epidemic and the challenges. Int. J. Antimicrob. Agents 55, 105924 (2020).

8. Yamamoto, N., Matsuyama, S., Hoshino, T. & Yamamoto, N. Nelfinavir inhibits replication of severe acute respiratory syndrome coronavirus 2 in vitro. http://biorxiv.org/lookup/doi/10.1101/2020.04.06.026476 (2020) doi:10.1101/2020.04.06.026476.

9. Casadevall, A. & Pirofski, L. The convalescent sera option for containing COVID-19. J. Clin. Invest. 130, 1545–1548 (2020).

10. Monteil, V. et al. Inhibition of SARS-CoV-2 infections in engineered human tissues using clinical-grade soluble human ACE2. Cell (2020).

11. Hoffmann, M. et al. SARS-CoV-2 Cell Entry Depends on ACE2 and TMPRSS2 and Is Blocked by a Clinically Proven Protease Inhibitor. Cell S0092867420302294 (2020) doi:10.1016/j.cell.2020.02.052.

12. Blanco-Melo, D. et al. SARS-CoV-2 launches a unique transcriptional signature from in vitro, ex vivo, and in vivo systems. http://biorxiv.org/lookup/doi/10.1101/2020.03.24.004655 (2020) doi:10.1101/2020.03.24.004655.

13. Maravillas-Montero, J. L. et al. Cutting edge: GPR35/CXCR8 is the receptor of the mucosal chemokine CXCL17. J. Immunol. Baltim. Md 1950 194, 29–33 (2015).

14. Thomas, A. C., Tattersall, D., Norgett, E. E., O’Toole, E. A. & Kelsell, D. P. Premature terminal differentiation and a reduction in specific proteases associated with loss of ABCA12 in Harlequin ichthyosis. Am. J. Pathol. 174, 970–978 (2009).

15. Wilk, E. et al. RNAseq expression analysis of resistant and susceptible mice after influenza A virus infection identifies novel genes associated with virus replication and important for host resistance to infection. BMC Genomics 16, 655 (2015).

16. Kanehisa, M., Sato, Y., Furumichi, M., Morishima, K. & Tanabe, M. New approach for understanding genome variations in KEGG. Nucleic Acids Res. 47, D590–D595 (2019).

17. Schoggins, J. W. & Rice, C. M. Interferon-stimulated genes and their antiviral effector functions. Curr. Opin. Virol. 1, 519–525 (2011).

18. Gillespie, K. A., Mehta, K. P., Laimins, L. A. & Moody, C. A. Human Papillomaviruses Recruit Cellular DNA Repair and Homologous Recombination Factors to Viral Replication Centers. J. Virol. 86, 9520–9526 (2012).

19. Zhang, Y. et al. [Clinical and coagulation characteristics of 7 patients with critical COVID-2019 pneumonia and acro-ischemia]. Zhonghua Xue Ye Xue Za Zhi Zhonghua Xueyexue Zazhi 41, E006 (2020).

20. Thompson, M. R. et al. Interferon γ-inducible Protein (IFI) 16 Transcriptionally Regulates Type I Interferons and Other Interferon-stimulated Genes and Controls the Interferon Response to both DNA and RNA Viruses. J. Biol. Chem. 289, 23568–23581 (2014).

21. Chan, M. C. et al. Tropism and innate host responses of a novel avian influenza A H7N9 virus: an analysis of ex-vivo and in-vitro cultures of the human respiratory tract. Lancet Respir. Med. 1, 534–542 (2013).

22. Jamaluddin, M., Tian, B., Boldogh, I., Garofalo, R. P. & Brasier, A. R. Respiratory Syncytial Virus Infection Induces a Reactive Oxygen Species-MSK1-Phospho-Ser-276 RelA Pathway Required for Cytokine Expression. J. Virol. 83, 10605–10615 (2009).

23. Meng, W. & Takeichi, M. Adherens junction: molecular architecture and regulation. Cold Spring Harb. Perspect. Biol. 1, a002899 (2009).

24. Lin, L.-T. & Richardson, C. The Host Cell Receptors for Measles Virus and Their Interaction with the Viral Hemagglutinin (H) Protein. Viruses 8, 250 (2016).

25. Lambert, D. W., Clarke, N. E., Hooper, N. M. & Turner, A. J. Calmodulin interacts with angiotensin-converting enzyme-2 (ACE2) and inhibits shedding of its ectodomain. FEBS Lett. 582, 385–390 (2008).

26. Hein, M. Y. et al. A human interactome in three quantitative dimensions organized by stoichiometries and abundances. Cell 163, 712–723 (2015).

27. Bertram, S., Glowacka, I., Steffen, I., Kühl, A. & Pöhlmann, S. Novel insights into proteolytic cleavage of influenza virus hemagglutinin. Rev. Med. Virol. 20, 298–310 (2010).

28. Kam, Y.-W. et al. Cleavage of the SARS coronavirus spike glycoprotein by airway proteases enhances virus entry into human bronchial epithelial cells in vitro. PloS One 4, e7870 (2009).

29. Kawase, M., Shirato, K., van der Hoek, L., Taguchi, F. & Matsuyama, S. Simultaneous treatment of human bronchial epithelial cells with serine and cysteine protease inhibitors prevents severe acute respiratory syndrome coronavirus entry. J. Virol. 86, 6537–6545 (2012).

30. de Aberasturi, A. L. & Calvo, A. TMPRSS4: an emerging potential therapeutic target in cancer. Br. J. Cancer 112, 4–8 (2015).

31. Bugge, T. H., Antalis, T. M. & Wu, Q. Type II Transmembrane Serine Proteases. J. Biol. Chem. 284, 23177–23181 (2009).

32. Tomlins, S. A. Recurrent Fusion of TMPRSS2 and ETS Transcription Factor Genes in Prostate Cancer. Science 310, 644–648 (2005).

33. Tomlins, S. A. et al. Role of the TMPRSS2-ERG gene fusion in prostate cancer. Neoplasia N. Y. N 10, 177–188 (2008).

34. Meng, T. et al. The insert sequence in SARS-CoV-2 enhances spike protein cleavage by TMPRSS. http://biorxiv.org/lookup/doi/10.1101/2020.02.08.926006 (2020) doi:10.1101/2020.02.08.926006.

35. Glowacka, I. et al. Differential downregulation of ACE2 by the spike proteins of severe acute respiratory syndrome coronavirus and human coronavirus NL63. J. Virol. 84, 1198–1205 (2010).

36. Huang, J., Song, W., Huang, H. & Sun, Q. Pharmacological Therapeutics Targeting RNA-Dependent RNA Polymerase, Proteinase and Spike Protein: From Mechanistic Studies to Clinical Trials for COVID-19. J. Clin. Med. 9, 1131 (2020).

37. Imai, Y. et al. Angiotensin-converting enzyme 2 protects from severe acute lung failure. Nature 436, 112–116 (2005).

38. Wadman, M., Couzin-Frankel, J., Kaiser, J. & Matacic, C. A rampage through the body. Science 368, 356–360 (2020).

39. Fang, L., Karakiulakis, G. & Roth, M. Are patients with hypertension and diabetes mellitus at increased risk for COVID-19 infection? Lancet Respir. Med. 8, e21 (2020).

40. Vaduganathan, M. et al. Renin–Angiotensin–Aldosterone System Inhibitors in Patients with Covid-19. N. Engl. J. Med. 382, 1653–1659 (2020).

41. Staudt, M. R., Salit, J., Kaner, R. J., Hollmann, C. & Crystal, R. G. Altered lung biology of healthy never smokers following acute inhalation of E-cigarettes. Respir. Res. 19, 78 (2018).

42. Gentleman, R. C. et al. Bioconductor: open software development for computational biology and bioinformatics. Genome Biol. 5, R80 (2004).

43. Leek, J. T., Johnson, W. E., Parker, H. S., Jaffe, A. E. & Storey, J. D. The sva package for removing batch effects and other unwanted variation in high-throughput experiments. Bioinforma. Oxf. Engl. 28, 882–883 (2012).

44. Johnson, W. E., Li, C. & Rabinovic, A. Adjusting batch effects in microarray expression data using empirical Bayes methods. Biostat. Oxf. Engl. 8, 118–127 (2007).

45. Law, C. W., Chen, Y., Shi, W. & Smyth, G. K. voom: precision weights unlock linear model analysis tools for RNA-seq read counts. Genome Biol. 15, R29 (2014).

46. Galili, T. dendextend: an R package for visualizing, adjusting and comparing trees of hierarchical clustering. Bioinformatics 31, 3718–3720 (2015).

47. Benjamini, Y. & Hochberg, Y. Controlling the False Discovery Rate: A Practical and Powerful Approach to Multiple Testing. J. R. Stat. Soc. Ser. B Methodol. 57, 289–300 (1995).

48. Falcon, S. & Gentleman, R. Using GOstats to test gene lists for GO term association. Bioinformatics 23, 257–258 (2007).

49. Wickham, H. Ggplot2: elegant graphics for data analysis. (Springer, 2009).

50. Wruck, W. & Adjaye, J. Meta-analysis of human prefrontal cortex reveals activation of GFAP and decline of synaptic transmission in the aging brain. Acta Neuropathol. Commun. 8, 26 (2020).

51. Chatr-aryamontri, A. et al. The BioGRID interaction database: 2017 update. Nucleic Acids Res. 45, D369–D379 (2017).

52. Butts, C. T. network: a Package for Managing Relational Data in R. J. Stat. Softw. 24, (2008).

